# ER stress relief drives ß-cell proliferation

**DOI:** 10.1101/2024.09.06.611615

**Authors:** Stephanie Bourgeois, Annelore Van Mulders, Yves Heremans, Gunter Leuckx, Lien Willems, Sophie Coenen, Laure Degroote, Julie Pierreux, Daliya Kancheva, Isabelle Scheyltjens, Kiavash Movahedi, Françoise Carlotti, Eelco de Koning, Xiaoyan Yi, Chiara Vinci, Yue Tong, Miriam Cnop, Harry Heimberg, Nico De Leu, Willem Staels

## Abstract

Regenerating endogenous pancreatic ß-cells is a potentially curative yet currently elusive strategy for diabetes therapy. Mimicking the microenvironment of the developing pancreas and leveraging vascular signals that support pancreatic endocrinogenesis may promote ß-cell regeneration. We aimed to investigate whether recovery from experimental hypovascularization of the endocrine pancreas, achieved by modulating the transgenic production of a VEGF-A blocker in ß-cells, could trigger mouse ß-cell proliferation. Serendipitously, we found that transgene overexpression in ß-cells induces endoplasmic reticulum (ER) stress and that subsequent relief from this stress stimulates ß-cell proliferation independent of vessel recovery. Transient *GFP* overexpression *in vivo* and chemical induction of ER stress *in vitro* replicated this ß-cell cycling response. Our findings highlight the potential side effects of ER stress due to transgene overexpression in ß-cells and assert that ER stress relief serves as a potent regenerative stimulus.

## Introduction

Diabetes mellitus is a chronic disease caused by insulin deficiency. Current treatments focus on regulating blood glucose levels without addressing the underlying ß-cell defect. To cure diabetes, restoring and preserving a functional ß-cell mass sufficient for fine-tuned glucose control is necessary. The prospect of cell replenishment through endogenous ß-cell regeneration is enticing and could be achieved by stimulating the proliferation of existing ß-cells, inducing the redifferentiation of dedifferentiated ß-cells, or generating new ß-cells from non-ß-cells [1]. However, the intricate processes involved in these ß-cell regeneration modalities are not fully understood.

Most ß-cells are generated during fetal development and ß-cell expansion by proliferation peaks in early postnatal life. Thereafter, proliferation rates decline to less than 0.1-0.5% in the adult human pancreas and 1% in adult mice [2–6]. Self-duplication remains the primary mechanism for maintaining the postnatal ß-cell mass in mice [7]. Stimulating proliferation of remaining ß-cells could restore the functional ß-cell mass in people with diabetes. Therefore, unraveling the signaling pathways that underlie ß-cell proliferation, especially in adulthood, is of significant interest.

ß-cells rely on systemic and microenvironmental cues to adapt their function and numbers [8]. Pancreatic islets are highly vascularized mini-organs with extensive endothelial cell (EC)-ß-cell crosstalk. Beyond supplying nutrients and oxygen, ECs provide a vascular basement membrane to ß-cells and secrete growth factors vital for ß-cell differentiation, survival, and proliferation [9]. Vascular Endothelial Growth Factor A (VEGF-A) is a key signaling molecule in the EC-ß-cell crosstalk [10]. Within islets, ß-cells are the main source of VEGF-A, which, by binding to the VEGF receptor-2 (VEGFR2) on ECs, stimulates their recruitment and proliferation [10]. Precise control of VEGF-A signaling is crucial for maintaining islet vascular homeostasis and functional ß-cell mass [11]. Elevated VEGF-A levels in pancreatic islets cause excessive vascular permeability, inflammation, and ß-cell death [12, 13]. Subsequent normalization of VEGF-A levels induces recovery of islet morphology and ß-cell mass by self-duplication and is supported by macrophages recruited from the bone marrow [14].

Previously, we used transgenic RIP-rtTA;TetO-sFLT1 mice (see methods) to conditionally overexpress and secrete soluble VEGFR1 (sFLT1) in ß-cells, thereby antagonizing VEGF-A signaling. This model enabled us to investigate the impact of islet hypovascularization and hypoxia on ß-cell function and proliferation during injury, adulthood [15] and pregnancy [16]. Despite causing mildly impaired glucose control, intra-islet hypovascularization did not affect baseline or stimulated ß-cell proliferation rates.

Here, we set out to examine whether recovery from experimental islet hypovascularization promotes ß-cell proliferation. Serendipitously, we observed increased ß-cell proliferation upon cessation of transgenic *sFLT1* overexpression, independently of blood vessel regrowth. Single-cell RNA sequencing (scRNA-seq) of pancreatic islet cells revealed that transgenic *sFLT1* overexpression causes significant endoplasmic reticulum (ER) stress in ß-cells and that subsequent stress relief coincides with cell cycle activation. Transient *GFP* expression in mouse ß-cells or *in vitro* exposure to the ER stress inducer thapsigargin [17] confirmed this replicative response. Our findings caution for side effects caused by transgene overexpression in ß-cells in genetically engineered mouse models and offer insight into how ER stress impacts ß-cell proliferation.

## Materials and methods

### Transgenic mouse models

Rat insulin gene promoter (RIP)-reverse tetracycline-dependent transactivator (rtTA) mice (RIP-rtTA) [18, 19] and tetracycline operator (TetO)-*sFLT1* mice [20, 21], both on a mixed CD1 background, were intercrossed to create double transgenic (dTg) mice in which human sFLT1 – a splice variant of the VEGF receptor 1 that inhibits signaling by VEGF-A, VEGF-B, and placental growth factor [22] – is secreted by ß-cells when doxycycline (DOX) is administered. RIP-rtTA;TetO-GFP,-LacZ mice were generated by intercrossing RIP-rtTA mice with TetO-GFP-LacZ mice (strain #018913, Jackson Laboratory, US). All animal procedures were performed following national laws and institutional guidelines. Mice were housed under standardized conditions (10 h dark/14 h light) and fed standard diet ad libitum. Mice were 6-8 weeks old when used in experiments. Genotyping was performed on ear biopsies using primers listed in **table 1**. DOX (0.4 mg/mL, Sigma-Aldrich, US) and BrdU (0.8 mg/mL, Sigma-Aldrich, US) were administered through the drinking water for 14 and 7 days, respectively. Tail vein blood glucose level and body weight were measured between 10 and 12 am, with prior fasting for 2 hours.

**Table 1:**
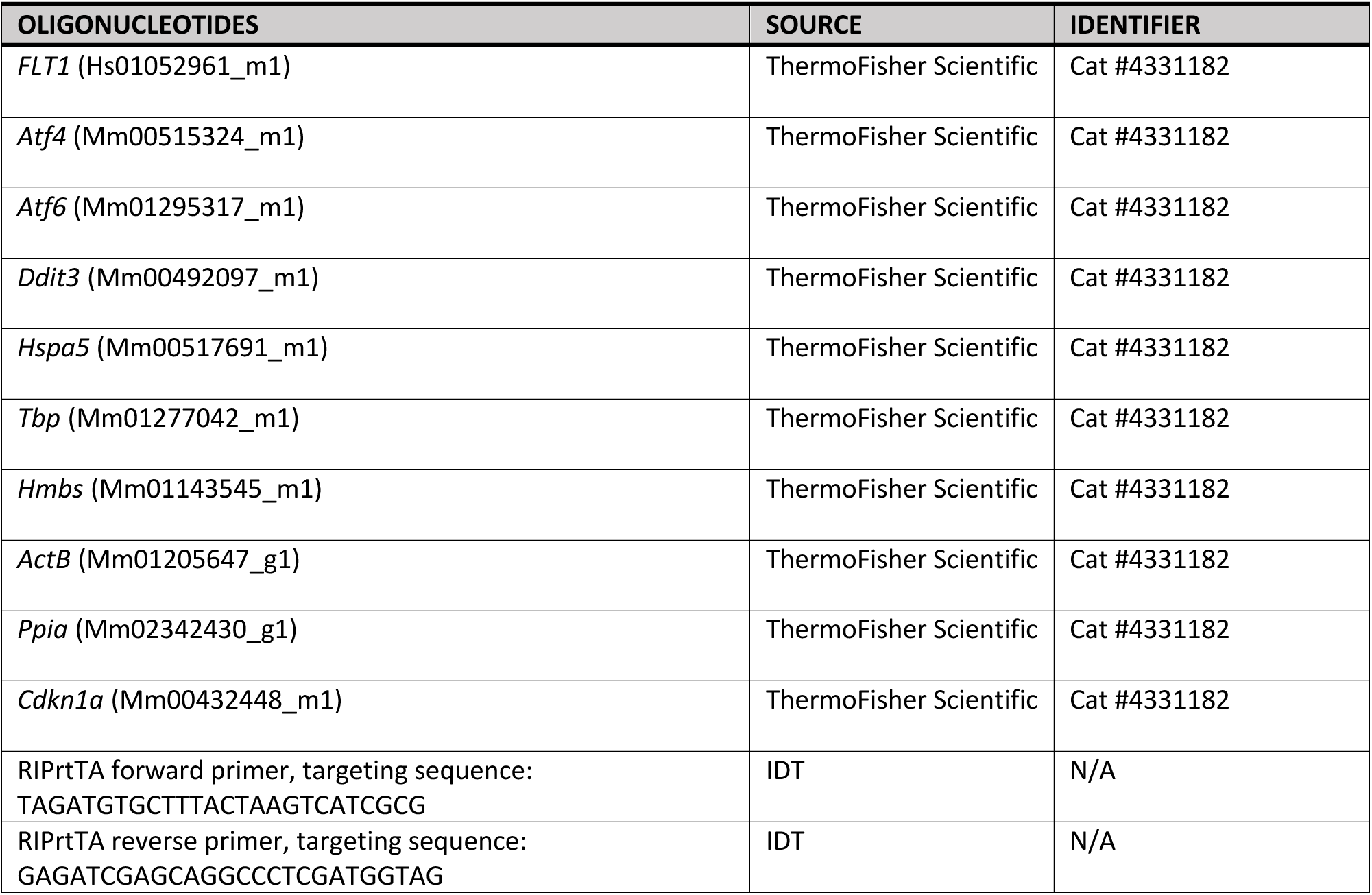
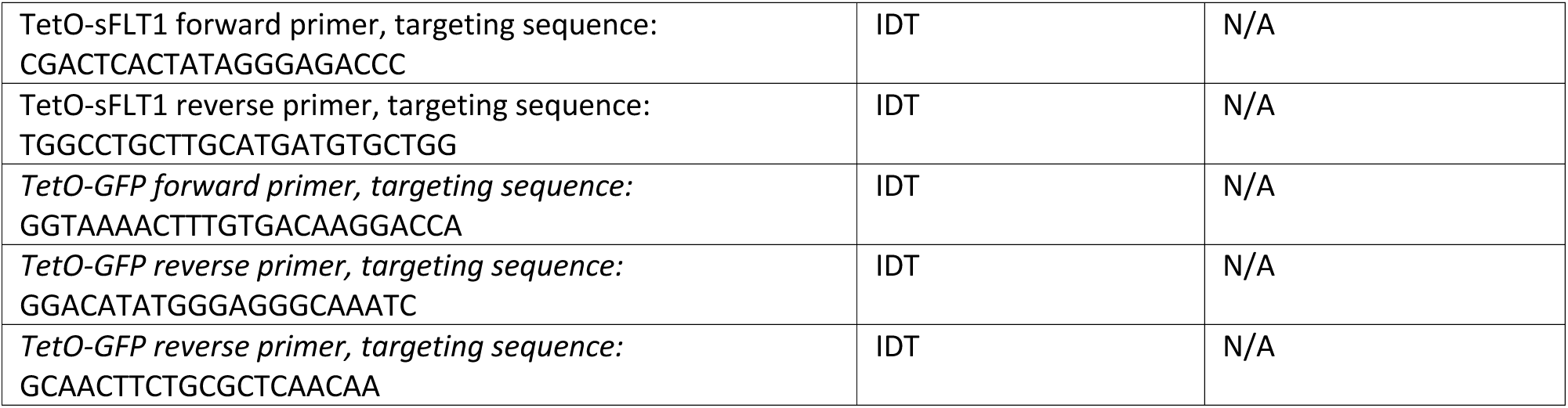
Primers used for gene expression analysis and genotyping.

### Protein analysis

Functional blood vessels were labeled by intravenous injection of biotinylated tomato lectin (Lycopersicon esculentum, Vector Laboratories, Burlingame, CA, USA). Five minutes after injection, blood cells were cleared from anesthetized mice with systemic PBS perfusion. Pancreas samples were fixed overnight in 10% (vol./vol.) neutral-buffered formalin (10% NBF) and then paraffin-embedded. Isolated islets were fixed for one hour in 10% NBF, entrapped in 2% (vol./vol.) agarose, and paraffin embedded.

For primary antibodies used, see **table 2**. An anti-FLT1 antibody was used to stain human sFLT1. Secondary antibodies were AlexaFluor-or Cyanine-labeled (Jackson ImmunoResearch, UK), and tomato lectin was visualized with AlexaFluor-labeled streptavidin (ThermoFisher Scientific, US). Nuclei were labeled with Hoechst 33342. Imaging was done using an Olympus BX61 microscope (Olympus, Japan) equipped with SmartCapture3 software (Digital Scientific UK, UK) or with a Zeiss LSM800 confocal microscope (Zeiss, Germany) equipped with ZEN software (Zeiss) and analyzed using Fiji software [23]. ß-cell proliferation was measured by BrdU or Ki67 positivity, and ß-cell volume was analyzed on at least 3% of the pancreas volume as previously described [24]. Proliferation in cultured islets was assessed with the Click-iT EdU kit (ThermoFisher Scientific, US).

**Table 2:** Primary antibodies used for protein expression analysis.

| ANTIBODY | SOURCE | IDENTIFIER |
| --- | --- | --- |
| Mouse monoclonal anti-BrdU | Dako | Cat#M0744; RRID: AB_10013660 |
| Rabbit monoclonal anti-CD81 | Cell Signaling Technology | Cat#10037; RRID: AB_2714207 |
| Rat monoclonal anti-F4/80 | Thermo Fisher Scientific | Cat#14-4801-85; RRID: AB_467559 |
| Mouse monoclonal anti-FLT1 | Santa Cruz | Cat#sc271789; RRID: AB_10710391 |
| Mouse monoclonal anti-Glucagon | Sigma-Aldrich | Cat#G2654; RRID:AB_259852 |
| Rabbit polyclonal anti-GLUT2 | Alpha Diagnostic | Cat#GT21-A; RRID: RRID: AB_1616640 |
| Guinea Pig anti-insulin | This paper | N/A |
| Rat monoclonal anti-Ki-67 | Thermo Fisher Scientific | Cat#14-5698-82; RRID: AB_10854564 |
| Rabbit polyclonal anti-MafA | Thermo Fisher Scientific | Cat #IHC-00352; RRID: AB_1279486 |
| Goat polyclonal anti-NKX6.1 | R&D Systems | Cat #AF5857, RRID: AB_1857045 |
| Rabbit monoclonal anti-p21 | Abcam | Cat #ab188224; RRID: AB_2734729 |
| Rabbit polyclonal anti-Phospho-Histone H3 (Ser10) | Cell Signaling Technology | Cat #9701; RRID: AB_331535 |
| Rat monoclonal anti-somatostatin | Abcam | Cat #ab30788, RRID: AB_778010 |
| Rabbit urocortin 3 | Huising et al. (PMID: 26076035) | N/A |

### Electron microscopy

Samples were fixed in 2.5% (vol/vol) glutaraldehyde and post-fixed in 1% (wt/vol) osmium-tetroxide and 2% (wt/vol) uranyl acetate, dehydrated in ethanol, immersed in propylene oxide (Honeywell Fluka, US), and embedded in Poly/bed 812 (Polysciences, Germany). 50-100 nm thin sections were cut with an Ultracut ultramicrotome (Reichert-Jung, US), mounted on copper grids, treated with uranyl acetate and lead citrate, and imaged with a Tecnai 10 transmission electron microscope (Philips, the Netherlands) using a mega viewG2 CCD camera (SIS-company, Germany).

### Mouse islet isolation, single-cell suspension preparation and *in vitro* assays

The main pancreatic duct was injected with 2 mL 1x HBSS (ThermoFisher Scientific, US) containing 0.140 mL CIzyme RI (PELOBiotech, Germany). The pancreas was transferred to a 50 mL tube containing 5 mL 1x HBSS supplemented with 0.25% (vol./vol.) BSA and 0.1 mg/mL kanamycin (kan) and digested for 17 minutes at 37°C with orbital shaking (95 strokes/min). The digestion was stopped by adding 30 mL ice-cold 1x HBSS supplemented with BSA/kan + 5% FBS. After centrifugation (1 min 1150 rpm), the pellet was resuspended in 10 mL 1x HBSS + BSA/kan + 5% FBS. The tissue was gently aspirated twice using a 14G needle fitted on a 20 mL syringe. Next, the digest was filtered over a 500-micron filter (pluriSelect Life Science, Germany) and transferred to a petri dish for handpicking of the islets using a stereomicroscope. For the single-cell RNA sequencing experiments, islets were dissociated into single cells with NeuroCult Dissociation Solution (STEMCELL Technologies, Canada). Cell number and viability were measured with a NucleoCounter NC-200 and Via1 cassettes (ChemoMetec, Denmark). Mouse primary islets were cultured at 37°C and 5% CO_2_ in RPMI 1640 (ThermoFisher Scientific, US), supplemented with 10% (vol./vol.) heat-inactivated FBS (Cytiva, US) and 100 U/mL penicillin/streptomycin (ThermoFisher Scientific, US). Isolated islets were exposed to varying concentrations of thapsigargin (Sigma-Aldrich, US) to induce ER stress. After 6 hours, brightfield images were taken using an EVOS M700 imaging system (ThermoFisher Scientific, US), and islets were collected in RLT buffer (Qiagen, Netherlands) with ß-mercaptoethanol and stored or transferred to new wells for washout before imaging and collection. RNA was isolated using RNeasy columns (Qiagen, Netherlands), and cDNA synthesis, RT, and qPCR were performed using mouse-specific Assays on demand[15] (ThermoFisher Scientific, US; **see Table 1**) with TaqMan Universal PCR Master Mix on a QuantStudio 12K Flex (ThermoFisher Scientific, US). A primer for *FLT1* was used to quantify human *sFLT1* expression. Reference genes were identified using geNorm analysis in qBase+ software[25] (Biogazelle, Belgium), and the geometric mean of multiple reference genes was used. EdU (10 μM, Click-iT® EdU Imaging Kits, ThermoFisher Scientific, US) was added to the medium for a 72-hour washout in samples for immunostaining.

### Single-cell RNA sequencing

GEMs and scRNA-seq libraries were prepared with the Chromium Next GEM Single Cell 3’ Kit v3.1, Chromium Next GEM Chip G Single Cell Kit, and the Dual Index Kit TT Set A (10x Genomics, US) according to the manufacturer’s instructions using the Chromium Controller (10x Genomics). GEM reverse-transcription (GEM-RT) was performed in a C1000 Touch Thermal Cycler (Bio-Rad, US) at 53°C for 45 min, 85°C for 5 min and ending at 4°C. GEM-RT cleanup was done using the Dynabeads MyOne SILANE reagent and the SPRIselect Reagent kit (Beckman Coulter, US). cDNA was amplified at 98°C for 3 min, 98°C for 15 sec, 63°C for 20 sec, 72°C for 1 min for a total of 11 to 12 cycles, followed by 1 min at 72°C and ending at 4°C. This was followed by enzymatic fragmentation, size selection, and adaptor and sample index attachment. The sequencing libraries were loaded onto a NovaSeq 6000 with the Novaseq S2 Reagent Kit (Illumina, US), and demultiplexing of the binary base call (BCL) files into individual samples was done using Cell Ranger mkfastq pipeline (v.6.0.2). The sFLT1 transgene sequence was integrated into the mouse reference genome (GRCm38, Ensembl release 98). Subsequently, mapping to this custom-modified genome, read filtering, barcode, and unique molecular identifier (UMI) counting were performed using kb-python v.0.25.1, a python wrapper of kallisto and bustools. Downstream analyses were conducted in R version 4.2.1. Low-quality cell barcodes were filtered out using emptyDrops from DropletUtils v.1.10.2 [26, 27]. Scater was then used to remove outlier cells based on UMI counts, gene number, and mitochondrial gene expression, as previously described [28]. For the mitochondrial gene percentage, a stricter cutoff of 20% was used instead of the Scater-defined threshold. Low-abundance genes were excluded as previously described[28], and doublet scores were assigned using scDblFinder (v.1.4.0) [29]. Next, the RNA expression matrix was further preprocessed by normalization, highly variable gene detection, scaling, and principal component analysis (PCA) using Seurat (v.3.2.3) [30]. Batch effect correction was applied using Harmony (v.1.0) [31] (θ-value = 0), considering only genes detected in both batches. Subsequent data analysis steps included Uniform Manifold Approximation and Projection (UMAP) and Louvain clustering on harmony-corrected PCA embeddings using the Seurat package [30]. Differential expression was analyzed with the Wilcoxon rank-sum test via Seurat’s “FindMarkers” function, with p-values adjusted using Bonferroni correction. GO analysis using Metascape (metascape.org) was conducted on differentially expressed genes with a Log2 fold change >0.25 and false discovery rate (FDR) <0.01. Significant GO Biological Processes terms had at least three associated genes, a p-value <0.01, and an enrichment score ≥1.5. To generate the ß-cell UMAP plots, the preprocessing steps were repeated on the ß-cell cluster to rectify any residual artifacts caused by doublets. We employed SCENIC v.1.2.4 [32] for gene regulatory network inference analysis, starting with raw, untransformed UMI counts as input data and following the recommended workflow. Genes with <30 UMI counts and expressed in <243 cells were filtered out. A co-expression network was established with GRNBoost2 (arboreto v.0.1.5), and the transcription factor-by-gene targets matrix was analyzed in R using SCENIC’s default settings. Regulon activity was computed using AUCell as previously described [32], and the results were visualized with Cytoscape (v.3.9.1) [33]. Trajectory analysis of the ß-cell subset was done using Monocle3 [34], with the root set at cluster 4 cells - predominantly found in the -DOX condition. The graph_test function visualized gene expression patterns by identifying modulated genes (q-value = 0 ΙMoran’s I score >0.35) along the trajectory.

### Quantification and statistical analysis

Data were statistically analyzed using (un)paired Student t-test, one-way ANOVA, two-way ANOVA or a mixed model for repeated measures analysis with Tukey post-hoc test (GraphPad Prism v.9.1.0, http://www.graphpad.com). The statistical details for each graph can be found in the corresponding figure legend. The number of animals or technical replicates used is represented as “n”. Results are presented as mean ± SEM, with p≤0.05 considered significant.

## Results

### Transient sFLT1 overexpression induces ß-cell proliferation independently of vessel recovery

To study the effect of intra-islet blood vessel ablation and regrowth on ß-cell proliferation, we used RIP-rtTA;TetO-sFLT1 mice. In this model, doxycycline (DOX) administration induces sFLT1 expression and secretion by ß-cells [15, 16], antagonizing vessel survival and angiogenesis by scavenging VEGF-A near ß-cells (**Fig. 1A**). The effects of transient sFLT1 expression on intra-islet vascularization and ß-cell proliferation were examined by administering DOX for 14 days followed by its withdrawal (WD) for 1, 4, or 7 days during which BrdU was administered (**Fig. 1B**). Double-transgenic (dTg) mice not receiving DOX and single-transgenic (sTg) TetO-sFLT1 mice that received DOX, referred to as -DOX and sTg mice, were used as controls. Islets isolated from +DOX mice contained high *sFLT1* transcript levels which decreased significantly after 4 days of DOX withdrawal (WD4D) (**Fig. 1C**). The expression of sFLT1 protein in ß-cells (+DOX: 87 ± 4.9%) and its decrease over time during WD were confirmed with immunostainings, however, at WD7D, half of the ß-cells (49 ± 4.8%) remained sFLT1-positive (**Fig. 1D,E**). Fasting glycemia was elevated in +DOX mice compared to sTg and -DOX mice but gradually normalized after WD (**Suppl. Fig. 1A**). The impact of transient sFLT1 expression on islet vascularization was assessed by quantification of the area, size, and number of islet vessels. The intra-islet blood vessel area (% lectin/insulin area) was strongly reduced in +DOX compared to sTg and -DOX controls. Surprisingly, at WD7D islet vascularization did not recover (**Fig. 1F,G** and **Suppl. Fig. 1B,C**). Even until WD 6 weeks, a subset of ß-cells remained sFLT1-positive, and islet hypovascularization persisted (**Suppl. Fig. 1D-F**). Intriguingly, cumulative BrdU labeling showed a nearly 3-fold increase in ß-cell proliferation in WD7D (14 ± 1.3%) compared to -DOX (5 ± 0.6%) and sTg (4 ± 0.2%) (**Fig. 1H,I**). The fraction of Ki67 and phospho-histone H3 (pHH3)-positive ß-cells peaked on WD4D (**Fig. 1J-L**) and insulin volume at WD7D was increased compared to controls (**Fig. 1M**). As transient islet hypervascularization following ß-cell-specific VEGF-A overexpression stimulates ß-cell proliferation following active recruitment of bone marrow-derived macrophages [14], we quantified intra- and peri-islet macrophages in our model. However, we found no differences between the experimental groups (**Suppl. Fig. 1G-I**). In summary, sFLT1 overexpression in ß-cells significantly reduces intra-islet blood vessels persisting at least 6 weeks after cessation of transgene overexpression. ß-cell cycling increases after transient *sFLT1* overexpression without significant recovery of islet ECs or changes in macrophage numbers.

**Figure 1:**
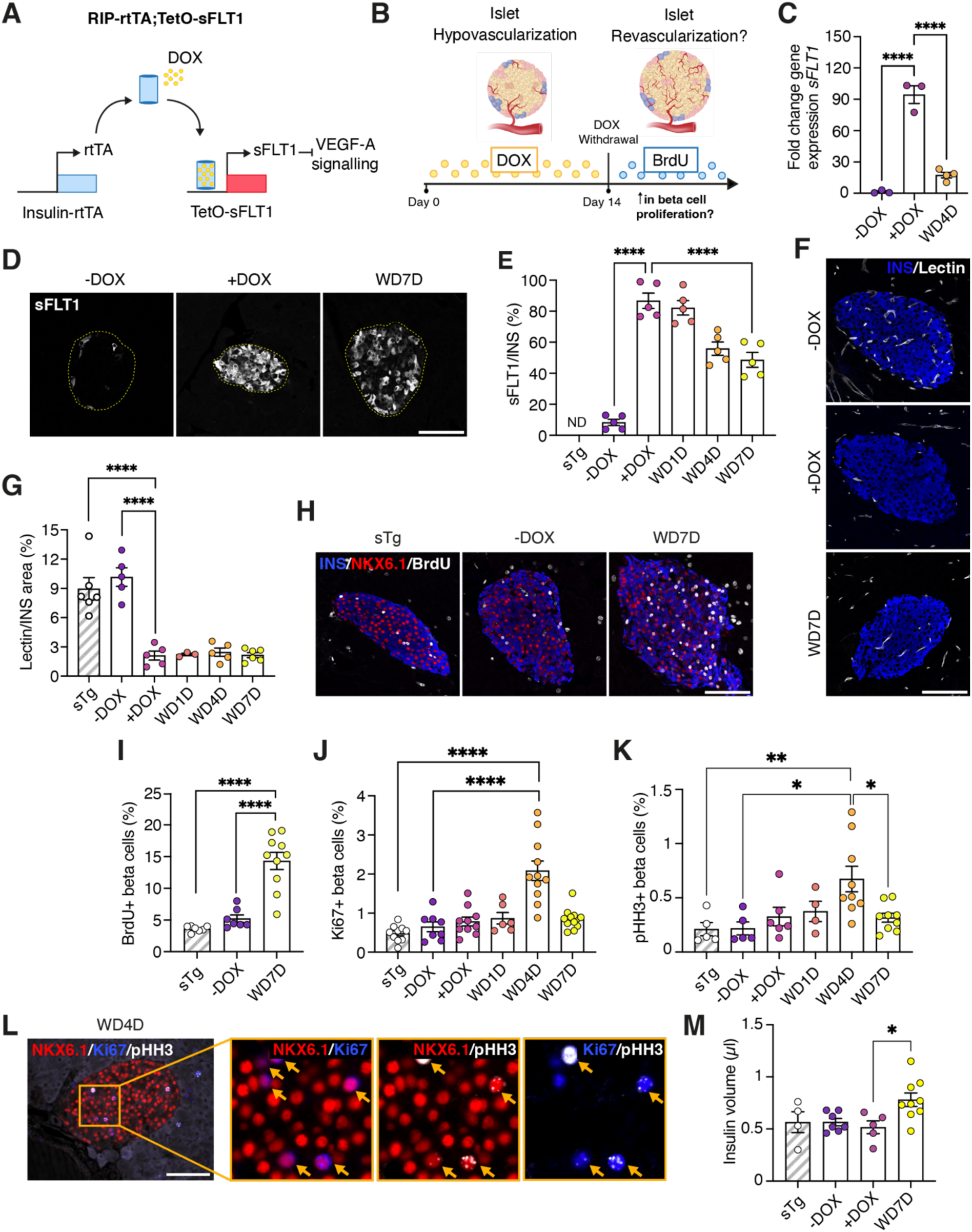
Transient sFLT1 overexpression induces ß-cell proliferation independently of vessel recovery. **(A)** Schematic of the RIP-rtTA;TetO-sFLT1 transgenic mouse model. **(B)** Experimental approach to study the effects of transient islet hypovascularization. **(C)** sFLT1 expression (fold change to -DOX) in pancreatic islets isolated from RIP-rtTA;TetO-sFLT1 mice, normalized to the geometric mean of ActB and Ppia. **(D)** Pancreatic sections immunostained for sFLT1. **(E)** sFLT1/INS area per pancreas. **(F)** Pancreatic sections immunostained for INS and lectin (functional blood vessels). **(G)** Lectin/INS area per pancreas. **(H)**Pancreatic sections immunostained for INS, NKX6.1 (ß-cell marker), and BrdU. **(I)** ß-cell cycling based on cumulative BrdU labeling for 7 days. **(J)** ß-cell proliferation based on Ki67-positivity. **(K)** Percentage of ß-cells in mitosis based on pHH3-positivity. **(L)** Immunostaining for Ki67, NKX6.1, and pHH3 on pancreatic sections. The arrows indicate proliferating ß-cells. **(M)** Insulin volume per pancreas. Data are mean ± SEM. One-way ANOVA followed by Tukey’s post-hoc test (Fig. 1C,E,G,I,J,K,M). *p ≤ 0.05, **p ≤ 0.01, ****p ≤ 0.0001. Scale bar = 100 µm. (rtTA = reverse tetracycline-controlled transactivator, sTg = single-transgenic, DOX = doxycycline, WD = withdrawal).

### sFLT1 overexpression causes ER stress and loss of end-stage maturation markers in ß-cells

To investigate the islet cell response to hypovascularization and the pathways driving ß-cell proliferation, we performed scRNA-seq on -DOX (8,699 cells), +DOX (19,135 cells), and +DOX WD4D (10,528 cells) islet cells. We annotated ß-, α-, δ-, and PP-cells, ECs, fibroblasts, acinar and ductal cells, T- and B-cells, macrophages, and dendritic cells based on marker gene expression (**Fig. 2A,B**). *sFLT1* transgene RNA overexpression, identified by “*pFUSE-1*” and “*hsFLT1*” expression, was detected in +DOX ß-cells (**Fig. 2C, Suppl. Fig. 2A,B** and Star Methods). Next, we analyzed differential gene expression between the different conditions. A hypoxia response was evident in +DOX ß-, α-, and δ-cells, as indicated by the upregulation of *Vegfa* and genes involved in glycolysis (**Suppl. Fig. 2C**). Upregulated genes in +DOX ß-cells encompassed ER stress and unfolded protein response (UPR) genes, including *Sdf2l1* [35]*, Pdia6* [36]*, Dnajc3* [37], *Fkbp11* [38]*, Atf4, Ddit3, Hspa5*, *Atf5* [39]*, and Nupr1* [40] (**Fig. 2C,D**). *Cdkn1a*, encoding p21 and *Chgb* (chromogranin B), a protein involved in trafficking secretory proteins into budding insulin granules [41], were also upregulated (**Fig. 2C**), alongside *Cd81* and *Aldh1a3,* markers of immaturity, dedifferentiation and dysfunction [42, 43] (**Fig. 2C**). Additionally, several identity and end-stage maturation markers were decreased in +DOX ß-cells, including *Ins1*, *Mafa*, *Slc2a2*, *Ucn3*, *Glp1r*, *Sytl4*, *Ppp1r1a*, and *Trpm5* [44] (**Fig. 2C, Suppl. Fig. 3A**).

**Figure 2:**
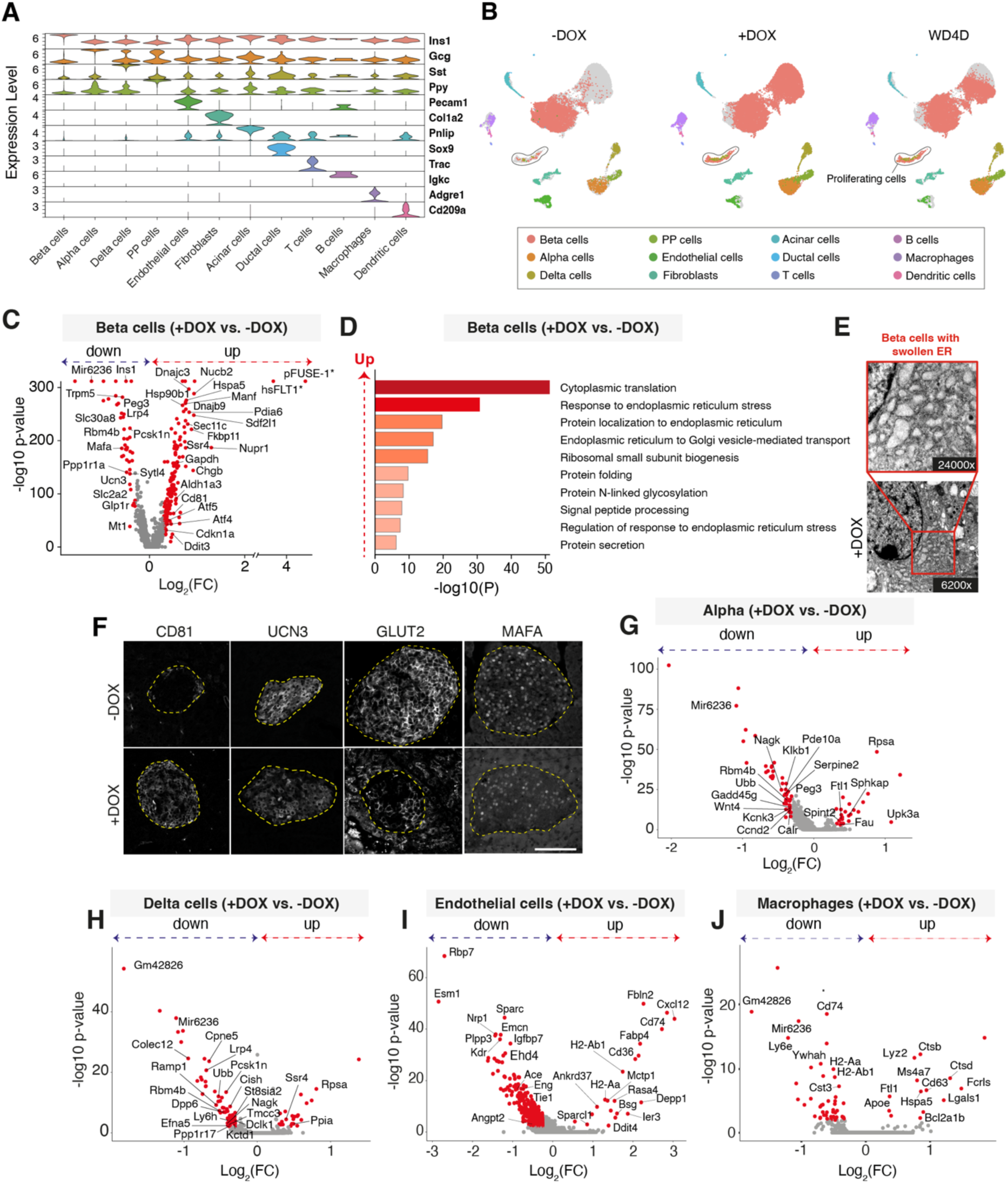
sFLT1 overexpression causes ER stress and loss of end-stage maturation markers in ß-cells. **(A)** Violin plot showing marker gene expression levels within each cluster of the islet cell scRNA-seq data.**(B)** UMAP plots of -DOX (8,699 cells), +DOX (19,135 cells), and WD4D (10,528 cells) islet cells (n = 2/condition). **(C)** Volcano plot displaying up- and downregulated genes in +DOX versus -DOX ß-cells. Genes with an absolute avg_log2FC > 0.3 and FDR < 0.01 are marked in red. **(D)** Gene ontology analysis of the significantly upregulated genes between -DOX and +DOX ß-cells. **(E)** Representative electron micrograph of +DOX ß-cells. **(F)** Pancreatic sections immunostained for CD81, UCN3, GLUT2, and MAFA in -DOX and +DOX. **(G-J)** Volcano plot displaying up- and downregulated genes in +DOX versus -DOX α-cells**(G)**, δ-cells **(H)**, endothelial cells **(I)**, and macrophages **(J)**. Genes with an absolute avg_log2FC expression value > 0.3 and FDR < 0.01 are marked in red. pFUSE-1 and hsFLT1, marked with *, represent the sequences used to detect sFLT1 transgene expression. Scale bar = 100 µm. (DOX = doxycycline, ER = endoplasmic reticulum, PP = pancreatic polypeptide, WD = withdrawal).

We confirmed presence of ER stress in +DOX ß-cells by electron microscopy, which showed an aberrantly swollen ER (**Fig. 2E**). In pancreatic sections, we also confirmed loss of ß-cell end-stage maturation markers based on decreased MAFA, GLUT2, and urocortin 3 and increased CD81 immunostaining in ß-cells (**Fig. 2F**). Next, we investigated the DOX effects on the transcriptome of other islet cell types. α-cells in +DOX retained normal expression levels of glucagon *(Gcg)* and other identity and maturation markers such as *Mafb*, *Arx*, *Irx1*, and *Irx2* [44]. Differentially expressed genes between +DOX and -DOX α-cells included genes involved in ER and oxidative stress such as *Upk3a* [45], *Rbm4b*, *Ubb*, *Gadd45g*, and *Calr* (**Fig. 2G and Suppl. Fig. 3B**). δ-cells in +DOX condition similarly retained key identity markers such as somatostatin (*Sst*), *Hhex*, and *Rbp4* [44] and downregulated genes (*Efna5*, *St8sia2*, *Dclk1*, *Cpne5*, and *Ramp1*) involved in cell morphogenesis, and neuronal and central nervous system development (**Fig. 2H and Suppl. Fig. 3C**). The +DOX ECs upregulated genes involved in glycolysis (*Tpi1*, *Gapdh* and *Pgk1*), antigen processing and presentation via MHC class II (*Cxcl12*, *Fbln2*, *Cd36*, *Cd74*, *H2-Aa,* and *H2-Ab1*) and downregulated genes related to EC identity and function (*Ace*, *Angpt2*, *Kdr*, *Eng*, *Esm1,* and *Tie1*) (**Fig. 2I and Suppl. Fig. 2C,3D**). Lastly, +DOX macrophages upregulated genes involved in glycolysis (*Tpi1*, *Gapdh*, *Pgk1* and *Pgam1*), lysosomal degradation and apoptosis (*Bcl2a1b*, *Ctsb*, *Ctsd,* and *Lgals1*) and downregulated genes involved in antigen processing and presentation via MHC class II (**Fig. 2J and Suppl. Fig. 2C, 3E**). To summarize, ß-cell-specific overexpression of sFLT1 increases ER stress and UPR in ß-cells concomitant with loss of ß- and EC maturation markers and induces a hypoxia response in all islet cells.

**Figure 3:**
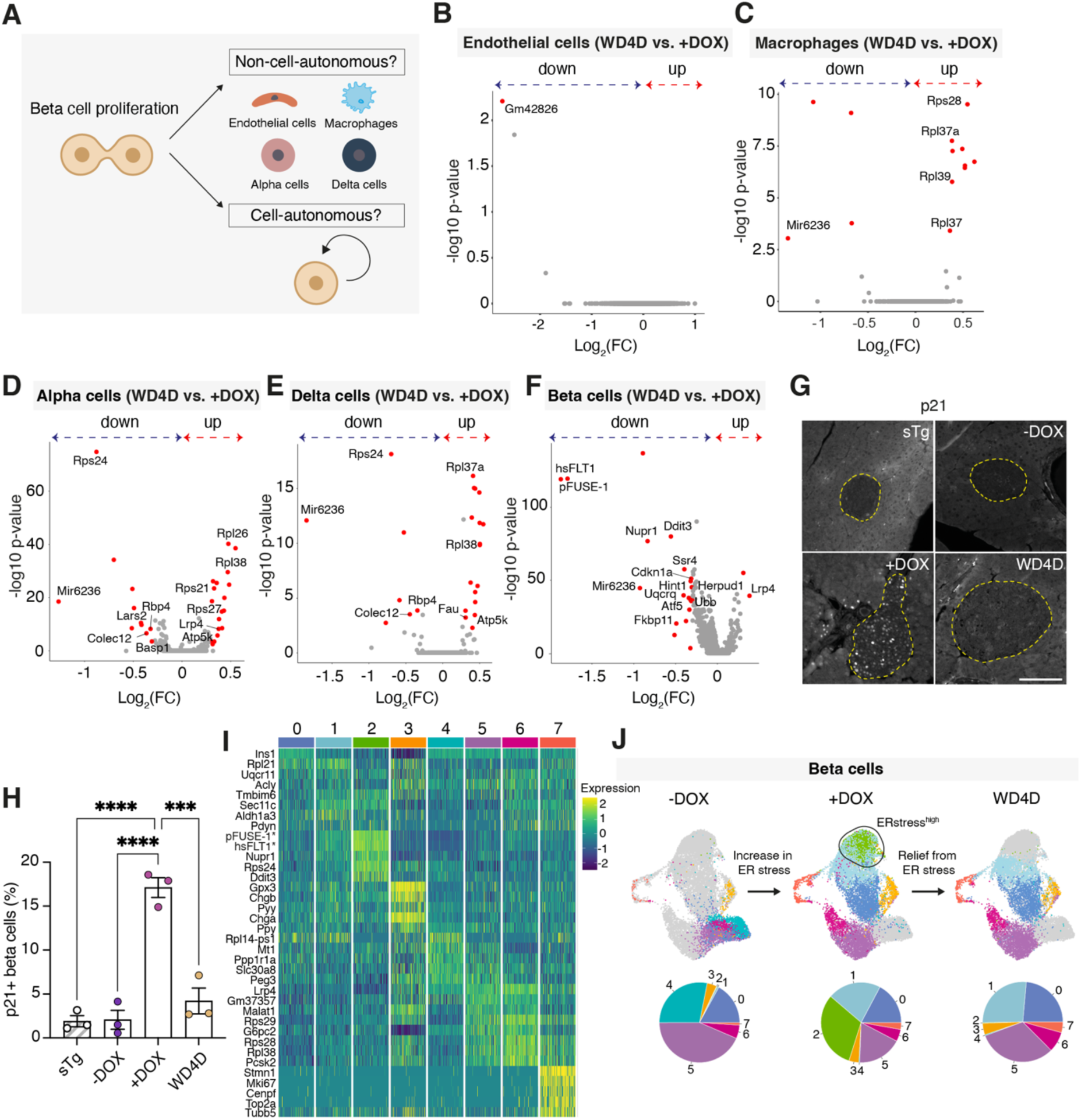
ER stress relief coincides with increased ß-cell proliferation. **(A)** Schematic model of the potential pathways driving ß-cell proliferation upon DOX withdrawal in RIP-rtTA;TetO-sFLT1 mice. **(B-F)** Volcano plot displaying up- and downregulated genes in WD4D versus +DOXendothelial cells **(B)**, macrophages **(C)**, α-cells **(D)**, δ-cells **(E)**, and ß-cells **(F)**. Genes with an absolute avg_log2FC > 0.3 and FDR < 0.01 are marked in red. **(G)** Pancreatic sections immunostained for P21. **(H)** Percentage of P21-positive ß-cells (sTg [n = 3], -DOX [n = 3], +DOX [n = 3], WD4D [n = 3]). **(I)** Heatmap representing the expression of top marker genes for each ß-cell cluster. **(J)** UMAP plots of 24,223 ß-cells coming from the -DOX, +DOX, and WD4D conditions. Pie charts indicating the percentages of each ß-cell cluster. Data are mean ± SEM. One-way ANOVA followed by Tukey’s post-hoc test (Fig. 3H). ***p ≤ 0.0001, ****p ≤ 0.0001. Scale bar = 100 µm. (sTg = single-transgenic, DOX = doxycycline, WD = withdrawal, ER = endoplasmic reticulum).

### Relief from ER stress coincides with increased ß-cell proliferation

To assess whether the observed increase in ß-cell proliferation after DOX WD (**Fig. 1I-L**) is due to cell-autonomous mechanisms or signals from other islet cell types (**Fig. 3A**), we performed scRNA-seq on islet cells and compared the transcriptomes of ECs, macrophages, α-, δ-, and ß-cells at the peak of ß-cell proliferation (WD4D) with those at +DOX. The scRNA-seq results confirmed the enrichment of proliferating ß-cells and the absence of EC recovery or macrophage recruitment in the WD4D condition (**Suppl. Fig. 2D-F**). Neither in ECs, macrophages, α- or δ-cells differential gene expression in potential ß-cell mitogens such as *Igf1*, *Fgf2*, *Hgf, Pdgf, Ctgf* or *Tgfb1*[1], was observed (**Fig. 3B-E**). In WD4D ß-cells, the *sFLT1* transgene, *Nupr1*, *Ddit3*, *Atf5, Fkbp11,* and *Cdkn1a* were significantly downregulated (**Fig. 3F**). Protein expression of p21, a negative cell cycle regulator, closely mirrored its gene expression profile with a significant increase in +DOX ß-cells and decrease in WD4D ß-cells (**Fig. 3G,H**). Most of the downregulated genes in WD4D ß-cells were involved in ER stress. qPCR for UPR marker genes on isolated islets from -DOX, +DOX, and WD4D confirmed ER stress induction and subsequent relief at the transcriptional level (**Suppl. Fig. 2G**). Szabat et al. reported that relief from basal ER stress upon knockout of the insulin genes induces ß-cell proliferation[46], we hypothesized that *sFLT1* overexpression caused ER stress in ß-cells and that ER stress relief upon DOX WD triggers ß-cell proliferation. To test this hypothesis, we re-clustered the ß-cell transcriptomes and identified 8 heterogenous ß-cell subsets (**Fig. 3I,J**). In addition, we used SCENIC [32] to identify potential transcription factors and their targets within each of the eight ß-cell subsets (**Fig. 4A**) for further characterization. Cluster 0 was present in all conditions and characterized by high *Ins1* expression and *Maz* and *Junb* regulon activity (**Fig. 3I,J and Fig. 4B**). In accordance with the loss of end-stage maturity markers (see above), cluster 1 - most abundant in +DOX and WD4D (**Fig. 3J**) – displayed relatively low *Pdx1* regulon activity and enriched activity of regulons associated with ER stress, such as Xbp1 (**Fig. 4B**). Furthermore, this cluster also showed higher expression of *Aldh1a3* (**Fig. 3I**). Absent in -DOX, cluster 2 appeared in +DOX and virtually disappeared in WD4D (**Fig. 3J**). These ß-cells exhibited the highest levels of the *sFLT1* transgene and ER stress genes *Nupr1* and *Ddit3*; hence, cluster 2 is referred to as the “ER stress^high^” cluster (**Fig. 3I,J**). This cluster showed highly enriched activity of regulons *Atf4*, *Cebpb*, *Cebpg*, and *Ddit3* (**Fig. 4C**), all associated with ER and cellular stress [39, 47, 48]*. Nupr1*, amongst the most robustly upregulated genes in +DOX ß-cells and amongst the most significantly downregulated genes in the WD4D condition, is a target of *Atf4*, *Cebpb*, *Cebpg,* and *Ddit3* (**Fig. 4C**), while *Cdkn1a* (another upregulated gene) is primarily regulated by *Cebpg*. Cluster 3, present in all conditions, had high *Gpx3, Chgb*, and *Chga* expression and was enriched in *Gata6*, *Sox4*, and *Foxp1* regulon activity; these transcription factors are interconnected through their roles in developmental processes, cell differentiation and disease states (**Fig. 3I,J and Fig. 4B**) [49–51]. Clusters 4 and 5 were predominant in the - DOX condition and likely represent more mature ß-cells, as indicated by the high expression of genes essential for ß-cell function such as *Ppp1r1a* and *Slc30a8*. These clusters were also characterized by high *Mafa* regulon activity (**Fig. 3I,J and Fig. 4B**). Cluster 6 exhibited enriched activity of *Creb3l2* and *Foxj2*, multifunctional transcription factors involved in stress responses, cell cycle regulation, and differentiation [52, 53]. Cluster 7 demarks proliferating ß-cells based on the expression of cell cycle genes (*Stmn1*, *Mki67*, *Cenpf*, *Top2a* and *Tubb5*) (**Fig. 3I**) and exhibited robust activity in gene regulatory networks associated with cell cycle transcription factors (**Fig. 4B**). To further dissect gene expression dynamics in the RIP-rtTA;TetO-sFLT1 ß-cells and their relationship to ß-cell cycling, we performed pseudotime trajectory analysis starting from ß-cells predominant in -DOX (as indicated in Fig. 3J). The top 78 genes exhibiting differential expression along the trajectory are highlighted (**Fig. 5A,B**). These genes include the transgene (*pFUSE-1* and *hsFLT1*), key regulators of ER stress and the UPR (*Nupr1*, *Ddit3*, *Pdia6*, and *Fkbp11*), as well as genes linked to proliferation (*Mki67* and *Cenpa*) (**Fig. 5C**). The trajectory showed a transient surge in ER stress and UPR activation preceding the upregulation of cell proliferation genes. Collectively, these data show that *sFLT1* overexpression by ß-cells under transcriptional control of the insulin promoter induces ER stress and that increased ß-cell proliferation coincides with ER stress relief in ß-cells corroborating our hypothesis that a transient surge in ER stress cell-autonomously promotes ß-cell cycling.

**Figure 4:**
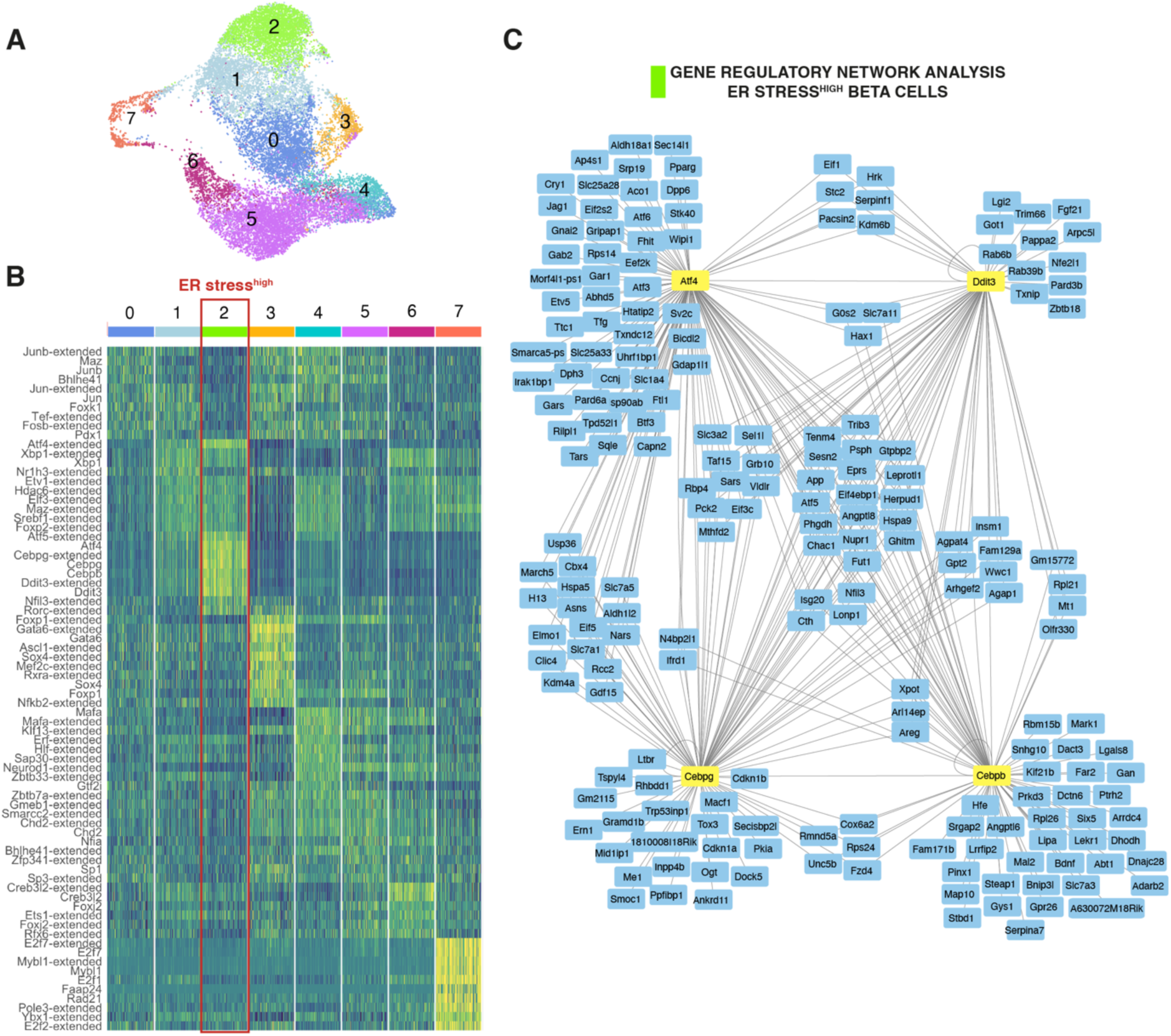
Gene regulatory network analysis of the ER stress^high^ ß-cell cluster reveals enriched Atf4, Ddit3, Cebpg, and Cebpb regulon activity. **(A)** UMAP showing the different ß-cell clusters in RIP-rtTA;TetO-sFLT1 mice. **(B)** Heatmap showing the top regulons with enriched activity within each ß-cell cluster. **(C)** Top regulons with enriched activity in the ER stress^high^ ß-cell cluster. The suffix “-extended” indicates low-confidence regulons (see STAR methods).

**Figure 5:**
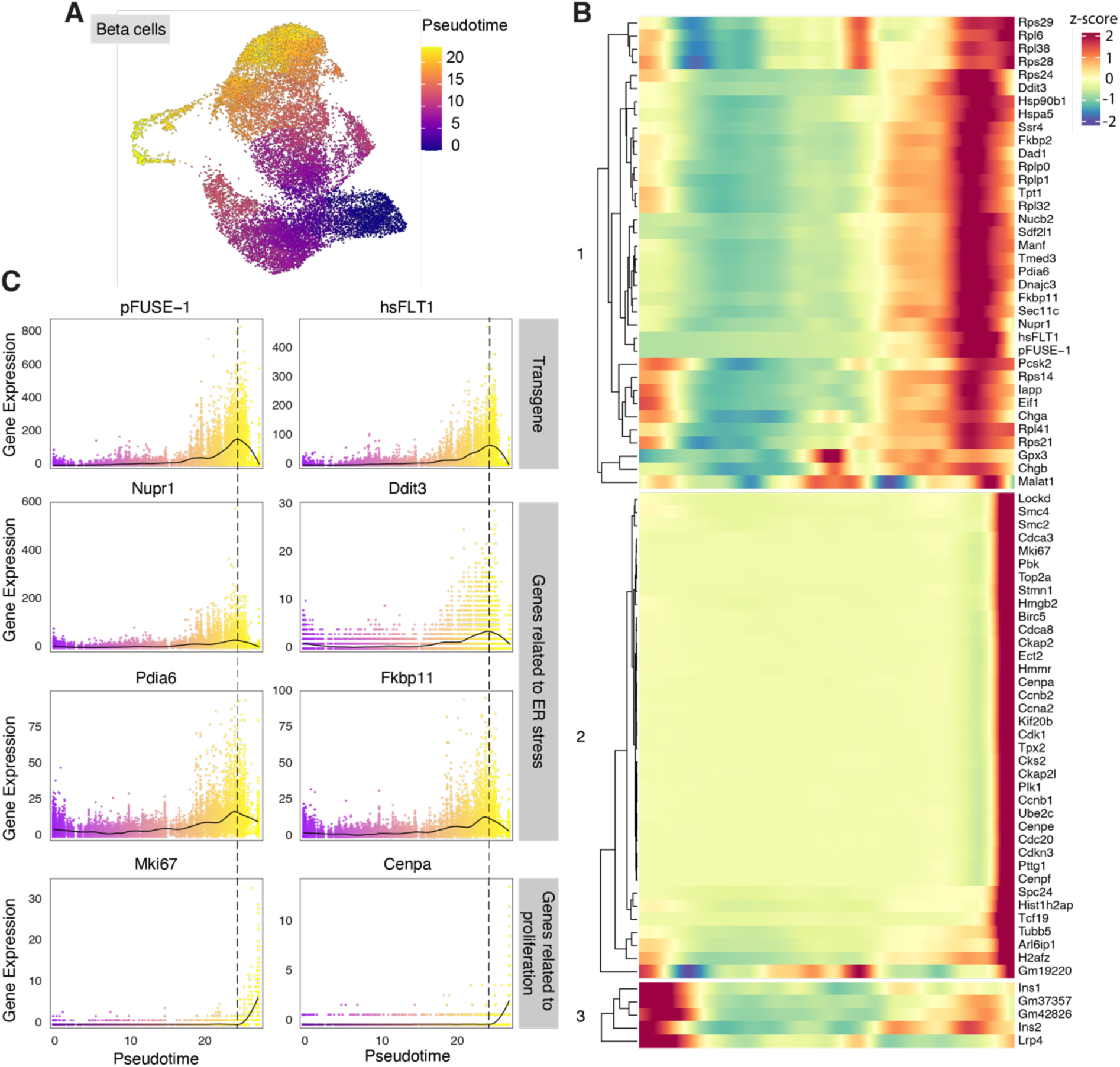
Pseudotime analysis shows high sFLT1 expression and ER stress occuring prior to increased ß-cell proliferation. **(A)** UMAP showing the pseudotime trajectory that was constructed from the 24,223 ß-cells derived from -DOX, +DOX, and WD4D RIP-rtTA;TetO-sFLT1 mice (n = 2/condition). -DOX ß-cells were defined as the root of the trajectory. Cells are colored by pseudotime. **(B)** Expression trends in pseudotime of the top 78 genes changing along the trajectory. Hierarchical clustering (using Euclidean distance) in three groups based on expression patterns along the pseudotime trajectory. The first group of genes are genes mostly related to protein synthesis, cellular stress responses, apoptosis regulation, and the transgene. The second group of genes are genes related to the regulation of the cell cycle and proliferation, and the third group of genes are involved in insulin production, non-coding sequences, and the Lrp4 receptor. The z-score for each gene’s expression value represents how much that value deviates from the average expression of that gene across the cells. **(C)** Expression pattern in pseudotime ordering of sFLT1 (indicated as pFUSE-1 and hsFLT1), Nupr1, Ddit3, Pdia6, Fkbp11, Mki67, and Cenpa.

### Conditional overexpression of GFP in ß-cells increases ß-cell proliferation

To test whether conditional overexpression of a random transgene under control of the insulin promoter induces ß-cell cycling, we developed RIP-rtTA;TetO-GFP transgenic mice in which DOX administration induced ß-cell-specific GFP expression (independently of islet vascularization) (**Fig. 6A**). A similar experimental design was used with DOX administration for 14 days, followed by DOX WD. Mice that did not receive DOX were used as controls. We validated *GFP* expression in +DOX islets and a significant decrease thereof at WD4D (**Fig. 6B**). Two hours fasting glycemia was measured weekly and showed no significant difference between the 2 groups (**Fig. 6C**). The abundance of ß-cell end-stage maturation markers MAFA and UCN3 did not differ on protein level between -DOX and +DOX mice expressing the GFP transgene (**Fig. 6D**). The UPR genes *Hspa5*, *Atf4*, *Ddit3,* and *Atf6* were significantly increased in +DOX and decreased in WD4D (**Fig. 6E**), similar to the sFLT1 model (**Suppl. Fig. 2G**). To expand on this comparison, we also assessed *Nupr1* and *Cdkn1a* gene expression. *Nupr1* expression was not modified, but *Cdkn1a* increased significantly in +DOX and decreased in WD4D (**Fig. 6E**). Notably, as in sFLT1 mice, transient GFP overexpression significantly increased ß-cell proliferation at WD7D (7 ± 0.5%) compared to -DOX (5 ± 0.5%) (**Fig. 6F,G**). In summary, conditional GFP overexpression by ß-cells transiently induces ER stress, with ß-cell proliferation increasing during the alleviation phase. These data suggest that any conditional transgene under the insulin promoter might evoke similar responses.

**Figure 6:**
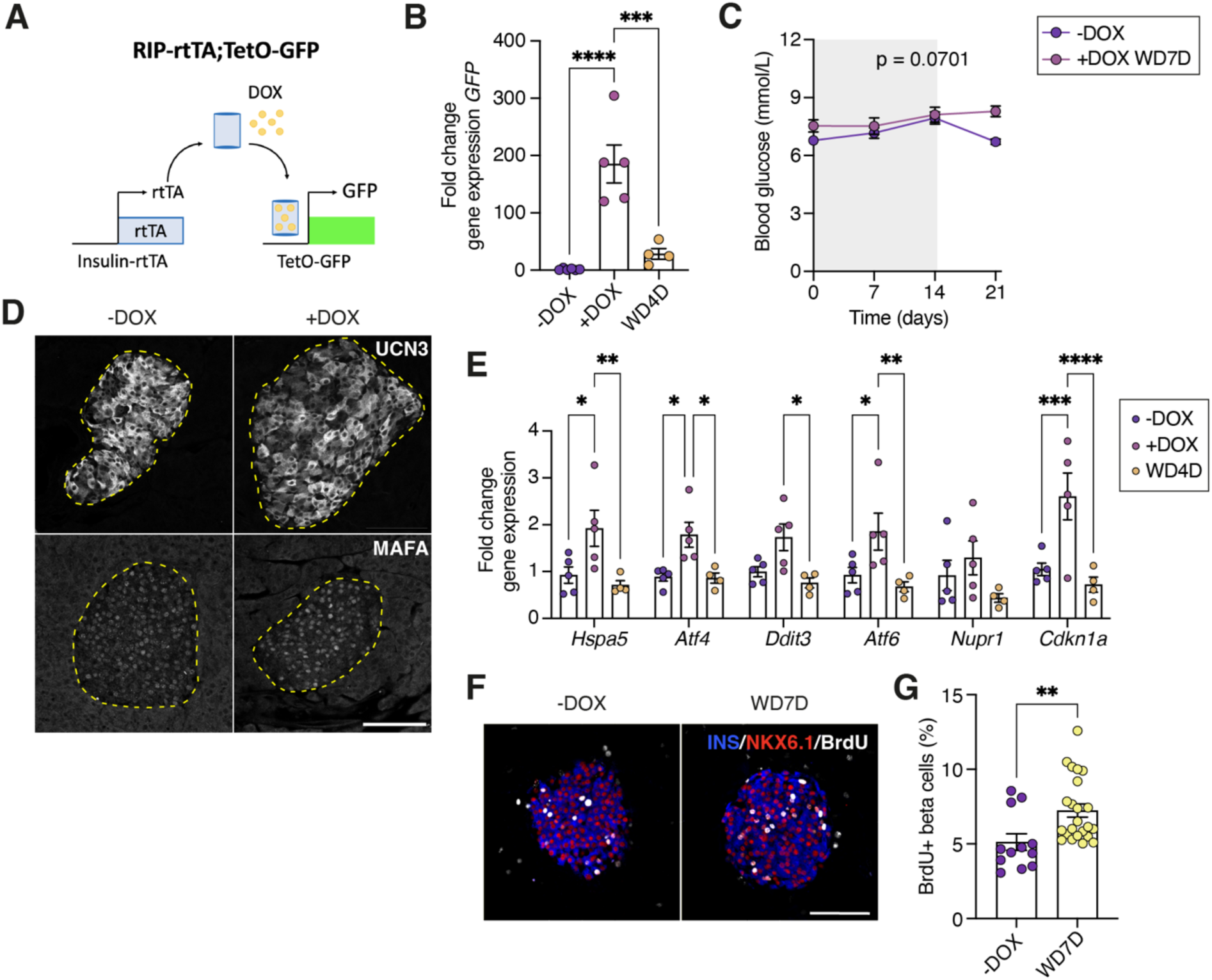
Conditional overexpression of GFP in ß-cells increases ß-cell proliferation. **(A)** Schematic of the RIP-rtTA;TetO-GFP transgenic mouse model. **(B)** GFP expression (fold change to -DOX) in pancreatic islets isolated from RIP-rtTA;TetO-GFP mice, normalized to the geometric mean of Tbp and Hmbs. (-DOX [n = 6], +DOX [n = 5], WD4D [n = 4]) **(C)** Two hours fasting glycemia (-DOX [n = 8], +DOX WD7D [n = 9]). The grey zone indicates the period during which DOX was administered. **(D)** Pancreatic sections immunostained for UCN3 and MAFA. **(E)** Hspa5, Atf4, Ddit3, Atf6, Nupr1, and Cdkn1a expression (fold change to -DOX) in pancreatic islets, normalized to the geometric mean of Tbp and Hmbs (-DOX [n = 5], +DOX [n = 5], WD4D [n = 4]). **(F)** Pancreatic sections immunostained for INS, NKX6.1, and BrdU. **(G)** Quantification of active ß-cell cycling after 7 days cumulative BrdU labeling (-DOX [n = 12], WD7D [n = 24]). Data are mean ± SEM. One-way ANOVA followed by Tukey’s post-hoc test (Fig. 6B,E). Mixed-effects analysis (Fig. 6C). Unpaired t-test (Fig. 6G). *p ≤ 0.05, **p ≤ 0.01, ***p ≤ 0.001, ****p < 0.0001. Scale bar = 100 µm. (rtTA = reverse tetracycline-controlled transactivator, DOX = doxycycline, WD = withdrawal)

### ER stress relief stimulates ß-cell proliferation in isolated mouse islets

To assess if transgene-independent relief from ER stress stimulates ß-cell proliferation, we exposed wildtype C57BL/6JRj islets *in vitro* to the ER stress inducer thapsigargin [17] (**Fig. 7A**). Based on the expression kinetics of *Ins1, Mafa, Slc2a2, Ucn3, Hspa5, Atf4, Ddit3,* and *Cdkn1a* in a dose-response assay (**Suppl. Fig. 4A**), we selected a concentration of 750 nM thapsigargin. We recapitulated ER stress induction and relief by 6 hours of thapsigargin exposure followed by a 24, 48, 72, or 96 hour washout period: thapsigargin induced ER stress with upregulation of *Ddit3*, *Atf4*, *Hspa5*, and *Cdkn1a* that normalized after 72 hours of washout (**Fig. 7B,C and Suppl. Fig. 4B**). Transient thapsigargin exposure did not induce cell death and kept the islet morphology intact (**Suppl. Fig. 4C**). ß-cell proliferation, as assessed by EdU incorporation and Ki67-positivity, was increased by nearly 3-fold after 72 hours of thapsigargin washout, an effect comparable to that of harmine treatment (**Fig. 7D-G**). These *in vitro* experiments further support our hypothesis that ER stress relief robustly stimulates mouse ß-cell proliferation.

**Figure 7:**
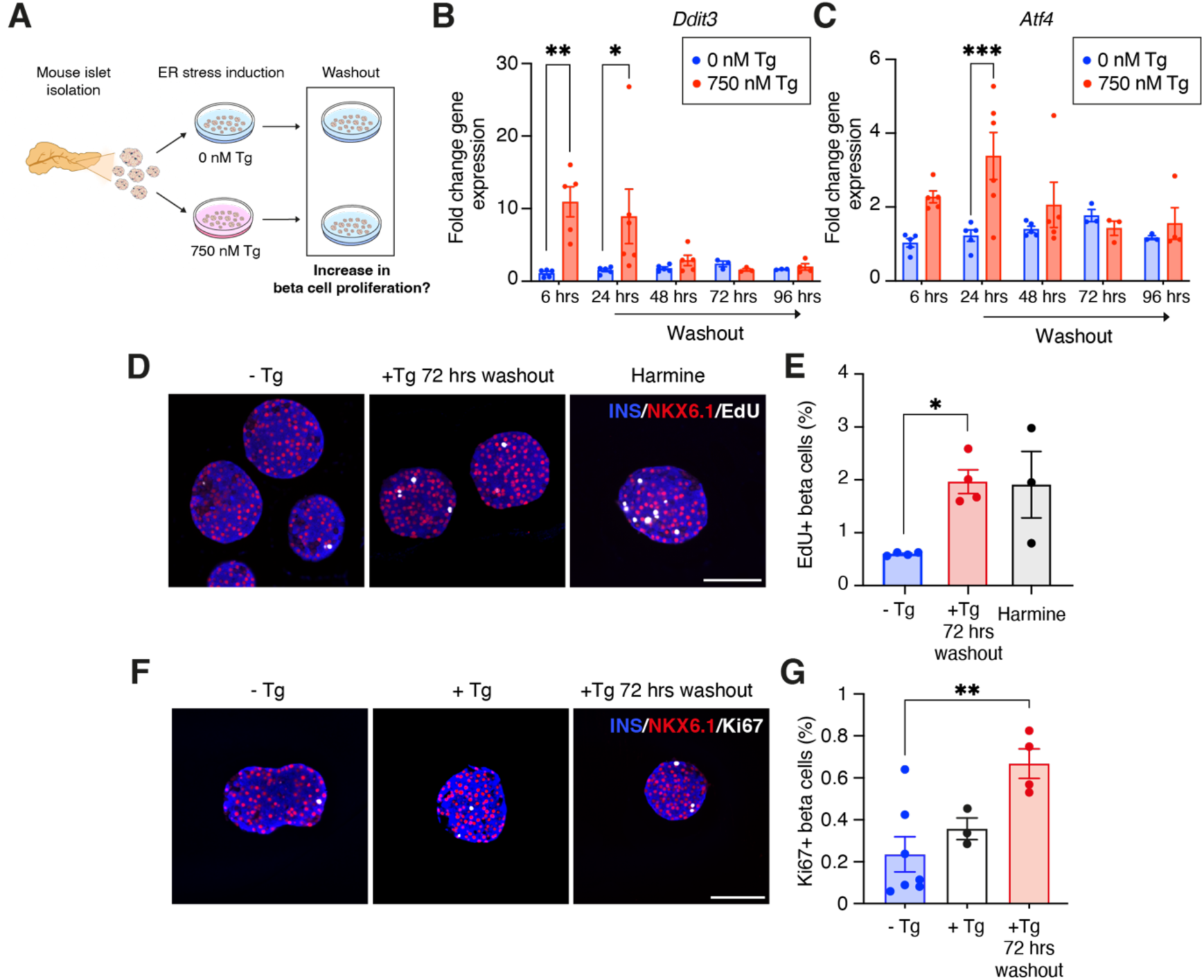
ER stress relief stimulates ß-cell proliferation in isolated mouse islets. **(A)** C57BL/6 mouse islets were exposed to 0 or 750 nM thapsigargin for 6 hours and ß-cell proliferation kinetics were examined during the washout period. Ddit3 **(B)** and Atf4 **(C)** expression (fold change to 6 hrs 0 nM Tg) normalized to the geometric mean of Ubc and Ppia. **(D)** Pancreatic islet sections immunostained for INS, NKX6.1 and EdU. **(E)** Quantification of active ß-cell proliferation after 72 hours of cumulative EdU labeling. **(F)** Pancreatic islet sections immunostained for INS, NKX6.1 and Ki67. **(G)** Quantification of active ß-cell cycling as assessed by Ki67-positivity. Data are mean ± SEM. Two-way ANOVA followed by Sidak’spost-hoc test (Fig. 7B,C), One-way ANOVA followed by Tukey’s post-hoc test (Fig. 7E,G). *p ≤ 0.05, **p ≤ 0.01, ***p < 0.001. Scale bar = 100 µm. (Tg = thapsigargin).

## Discussion

Finding effective methods to restore ß-cell mass is crucial to advance diabetes therapies. We hypothesized that mimicking vascular signals supportive of pancreatic endocrinogenesis during development could stimulate ß-cell generation during adulthood. Therefore, we set out to investigate the effects of intra-islet blood vessel ablation followed by revascularization. Transgenic overexpression of sFLT1 by ß-cells induced islet hypovascularization, and halting its overexpression increased ß-cell cycling, albeit in the absence of vessel recovery.

Brissova et al. showed that transient islet hypervascularization increased ß-cell proliferation through interactions among intra-islet macrophages, ECs, and extracellular matrix molecules [14, 54]. We found no significant difference, however, in macrophage numbers or their transcriptome between the different conditions in our model. Glucose, known for its mitogenic effects on ß-cells [1, 55, 56], was not the driver of this effect either, as ß-cell proliferation did not increase while fasting glycemia peaked but only did so afterward. This suggested a cell-autonomous process prompting the exploration of ß-cell-specific transcriptional changes.

Single-cell transcriptomics revealed reduced ß-cell identity markers, increased ER stress, and elevated UPR marker genes in *sFLT1-*overexpressing ß-cells. The observed loss of end-stage ß-cell maturation markers aligns with the need for EC-ß-cell crosstalk and nutrients to maintain functional maturity [57]. ER stress, exacerbated by transgene production and islet hypoxia [58], may have amplified this response [59, 60].

Using RIP-rtTA;TetO-GFP transgenic mice, we confirmed that also transient GFP overexpression in ß-cells induced ER stress followed by stimulation of ß-cell proliferation, though less pronounced than in RIP-rtTA;TetO-sFLT1 mice and without loss of canonical ß-cell markers’ expression. The difference in the mitogenic response is likely due to the hypoxic microenvironment in the sFLT1 model amplifying the ER stress [58] and pushing ß-cells into a more immature state. Unlike sFLT1, GFP is not a secreted protein. Despite this, we observed the induction of ER stress and its subsequent relief following GFP overexpression and its cessation. This observation aligns with findings in the mouse lens, where transgene overexpression induced ER stress and activated the UPR, regardless of whether the proteins were synthesized in the cytosol or ER [61]. Finally, *in vitro* modeling of transient ER stress by exposing wildtype mouse ß-cells to thapsigargin corroborated the mitogenic response to UPR relief.

The UPR is activated to prevent the accumulation of misfolded or unfolded proteins via three canonical branches: PERK/eIF2α, IRE1/XBP1, and ATF6. Upon BiP dissociation, these transducers elicit signaling that will halt protein translation, degrade misfolded proteins, and enhance protein folding [39]. Our results align with four studies that connect the UPR to ß-cell proliferation. First, Szabat et al. showed that baseline ER stress at physiological insulin production inhibits adult ß-cell proliferation. The loss of both *Ins1* and *Ins2* alleles stimulated ß-cell proliferation before the onset of hyperglycemia, coinciding with reduced expression of *Ddit3*, *Trib3*, *Atf4*, and *Atf5*, decreased *Xbp1* splicing, and increased eIF2α phosphorylation [46]. Second, Sharma et al. found that the UPR senses insulin demand and, via activation of the ATF6 branch, can promote ß-cell proliferation to increase ß-cell mass [62]. Third, Xin et al. performed scRNA-seq of cadaveric pancreatic islets from human donors and identified a INS^lo^UPR^high^ ß-cell population, suggested to consist of ß-cells recovering from high insulin production and enriched with proliferating ß-cells [63]. Fourth, the salt-inducible kinase (SIK) inhibitor HG-9-91-01 was found to induce ß-cell proliferation by a transient ATF6-dependent UPR. However, other signaling effectors downstream of SIK inhibition were required for increased ß-cell proliferation [64]. In summary, ß-cell proliferation and the UPR are closely linked. As noted by Szabat and colleagues [46], the specific pathways involved are likely context-dependent, with different branches of the UPR being activated under varying conditions.

Regenerative responses to ER stress relief are of major clinical interest as ER stress is closely related to ß-cell failure and the progression of type 1 and type 2 diabetes [65–67]. In mouse models of diabetes, the glucagon-like peptide 1 (GLP-1) agonist exendin-4 modulates ER stress responses, leading to improved ß-cell adaptation and survival [68]. Exendin-4 protects INS-1E cells and primary rat ß-cells, strengthening their defense mechanisms when exposed to the ER stressor cyclopiazonic acid [69]. GLP-1 agonists are now established as a first-line therapy for type 2 diabetes, and their beneficial effects may be partly attributed to the modulation of ER stress and the UPR. In new-onset type 1 diabetes, the initiation of insulin therapy after a variable period of increased metabolic workload due to decreased ß-cell numbers may replicate an ER stress/UPR response with subsequent relief and possibly contribute to the honeymoon phase observed in the weeks after starting insulin treatment. Given the regenerative potential of alleviating ER stress in our murine model systems, it is tempting to propose early intensive insulin therapy to reduce the strain on remaining ß-cells and promote their proliferation. In children with newly diagnosed type 1 diabetes, intensive diabetes management did not, however, prevent the decline in functional ß-cell mass [70, 71], and parenteral insulin therapy did not slow the progression from preclinical to clinical diabetes in the DPT-1 trial (NCT00004984). Nonetheless, it is worth exploring early insulin therapy to mitigate metabolic stress on remaining ß-cells in combination with immune interventions that halt the concurrent inflammatory stress.

## Conclusion

In summary, we demonstrate that a transient wave of ER stress can elicit ß-cell proliferation, highlighting unexpected and previously unknown effects of ER stress in commonly used transgenic mouse strains for ß-cell studies. In our study, the overexpression of *sFLT1* and *GFP* was driven by the strong Rat Insulin Promotor (RIP, or *Ins2),* which has previously been used to drive genes of interest in other ß-cell proliferation studies [14, 15, 72, 73]. Researchers should carefully consider evaluating ER stress in these transgenic strains, as it may bias interpretations of ß-cell behavior. Our findings that ER stress relief provides a regenerative impetus to ß-cells should encourage further studies on alleviating the metabolic workload of ß-cells in early-onset diabetes and could have potential implications for pharmacological intervention.

## Acknowledgements

We thank H. Azibou, N. Nasiri, M. Beya, A. Demarré, V. Laurysens and G. Stangé (Vrije Universiteit Brussel, Brussels, Belgium) for their valuable help.

## CRediT authorship contribution statement

Conceptualization: N.D.L., W.S. and H.H.; Investigation: S.B., W.S., A.V.M., L.W., S.C., J.P., L.D., Y.H., G.L., C.V., Y.T. and. I.S.; Software: G.L., S.B., D.K., and X.Y.; Resources: K.M., I.S., D.K., M.C., X.Y., F.C. and E.d.K.; Writing – Original Draft: S.B.; Writing – Review & Editing: W.S., N.D.L., Y.H., M.C. and H.H.; Funding Acquisition: H.H., N.D.L., and W.S.; Supervision: W.S., N.D.L, and H.H.

## Data and code availability

The single-cell RNA-sequencing data generated in this study are deposited at GEO (NCBI) with accession code GSE274443. Other data that support the findings of this study are available from the corresponding author upon request. There are no restrictions on data availability.

