## Supplementary data for "ER stress relief drives ß-cell proliferation"

**Supplementary Figure 1: (A)** Two hours fasting glycemia (sTg [n = 15], -DOX [n = 13], +DOX WD [n = 14]). The grey zone indicates the period during which DOX was administered. Quantification of the lectin-positive vessel size **(B)**, and number of lectin-positive vessels per insulin area **(C)**. **(D)** Pancreatic sections immunostained for sFLT1, insulin and lectin. **(E)** sFLT1/INS area per pancreas. **(F)** Lectin/INS area per pancreas. **(G)** Pancreatic sections immunostained for F4/80 and insulin. **(H)** Number of macrophages per $mm^2$ insulin area. **(I)** Number of intra-islet macrophages and of macrophages within a perimeter of 20 microns outside of the insulin area per $mm^2$ area. Data are mean ± SEM. Mixed-effects analysis (Suppl. fig. 1A). One-way ANOVA followed by Tukey's post-hoc test (Suppl. Fig. 1B,C,E,F,H,I). (sTg = single-transgenic, DOX = doxycycline, WD = withdrawal, WD2W = withdrawal 2 weeks WD4W = withdrawal 4 weeks, WD6W = withdrawal 6 weeks, MΦ = macrophage).

**Supplementary Figure 2:** UMAP plots showing the expression of the sFLT1 transgene, marked as hsFLT1 **(A)** and pFUSE-1 **(B)**. **(C)** Dot plots showing the expression of hypoxia marker genes in α-, ß-, δ-cells, macrophages and endothelial cells from -DOX, +DOX, and WD4D mice. Barplots showing the percentage of Proliferating ß-cells **(D)**, endothelial cells **(E)**, and macrophages **(F)** in isolated islets from -DOX, +DOX, and WD4D mice. **(G)** *Atf4*, *Atf6*, *Ddit3*, *Hspa5* and *Calr* gene expression (fold change to -DOX) in pancreatic islets isolated from RIP-rtTA;TetO-sFLT1 mice, normalized to the geometric mean of *ActB* and *Ppia.* Data are mean ± SEM. One-way ANOVA followed by Tukey's post-hoc test (Suppl. Fig. 2G). *p ≤ 0.05, **p ≤ 0.01. (Expr = expression, Avg = Average, DOX = doxycycline)

**Supplementary Figure 3:** Gene ontology analysis on differentially expressed genes between -DOX and +DOX ß-cells **(A)**, α-cells **(B)**, δ-cells **(C)**, endothelial cells **(D)**, and macrophages **(E)**, (FDR < 0.01 and abs($log_2$FC > 0.25). (DOX = doxycycline).

**Supplementary Figure 4: (A)** *Ins1, Mafa, Slc2a2, Ucn3, Hspa5, Atf4, Ddit3,* and *Cdkn1a* expression (fold change to 0 nM Tg) in pancreatic islets exposed to an increasing concentration of Tg for 6 hours, normalized to the geometric mean of *Ppia* and *Ubc*. **(B)** *Hspa5*, *Cdkn1a*, and *Ins1* expression (fold change to 0 nM Tg) in pancreatic islets exposed to either 0 or 750 nM Tg for 6 hours, followed by a washout period of 24, 48, 72, or 96 hours, normalized to the geometric mean of *Ppia* and *Ubc*. **(C)** Morphology of islets exposed for 6 hours to 0 or 750 nM Tg and of islets exposed for 6 hours to 750 nM Tg followed by a washout period of 72 hours. Scale bar = 650 µm. (Tg = thapsigargin).

**Supplementary figure 1**

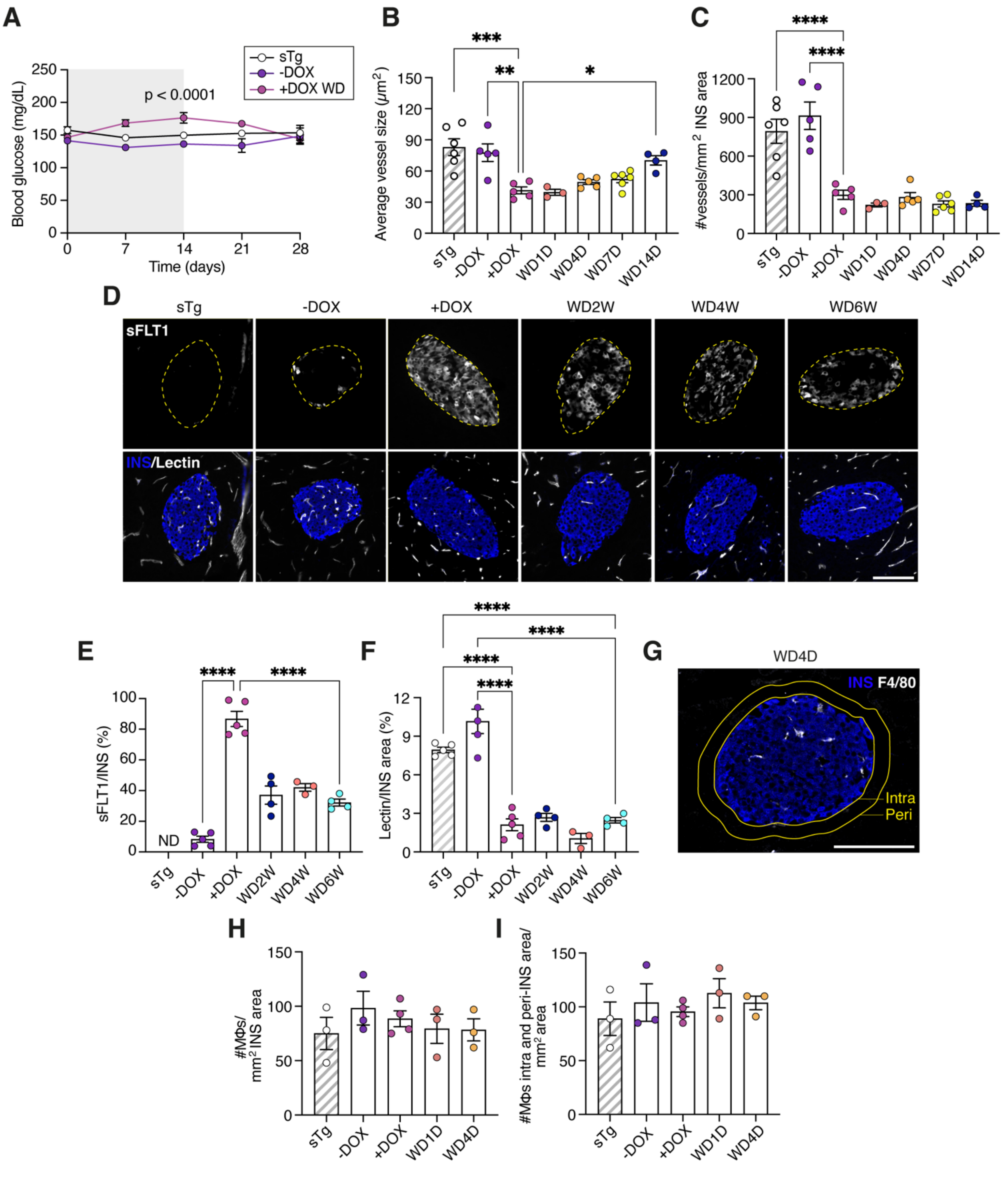

### Supplementary figure 2

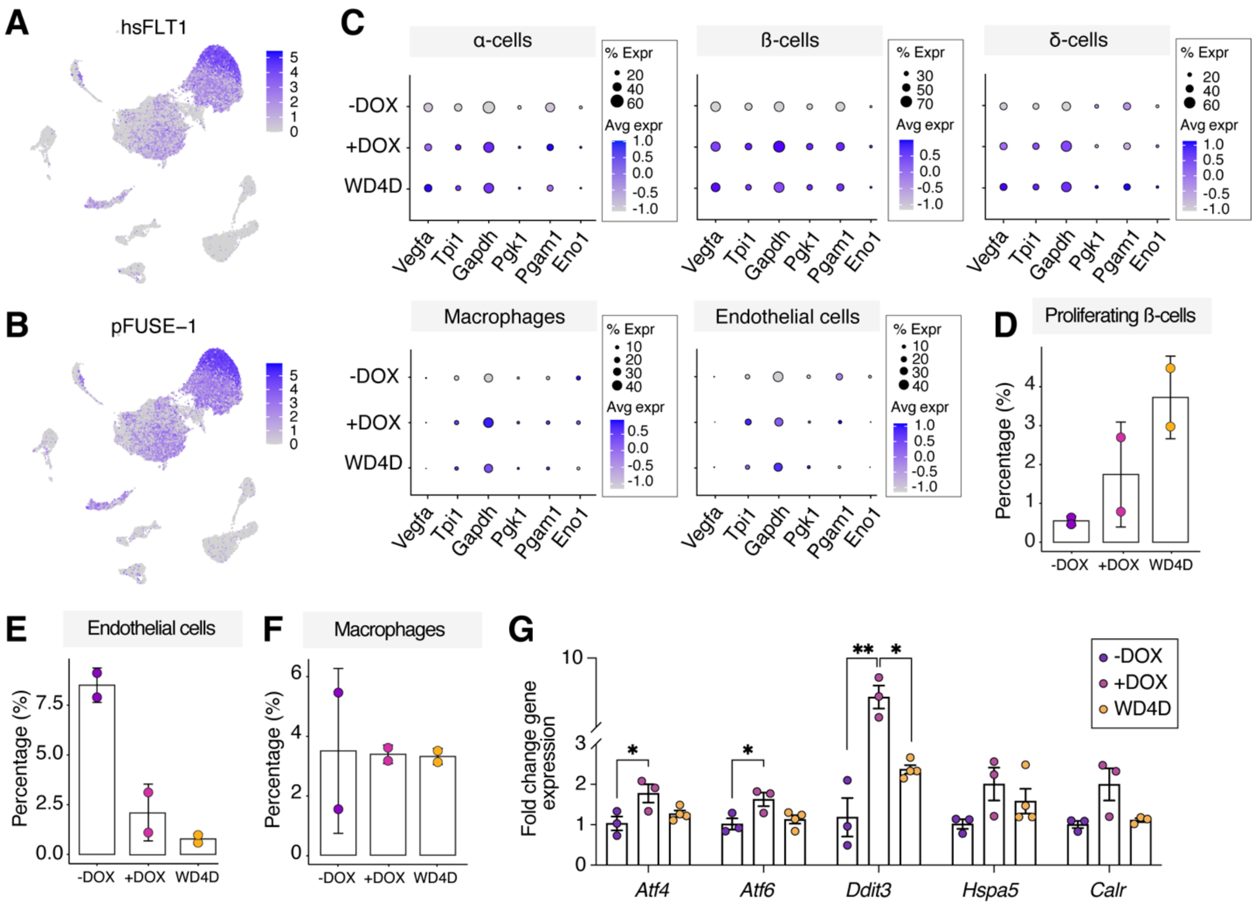

**Supplementary figure 3**

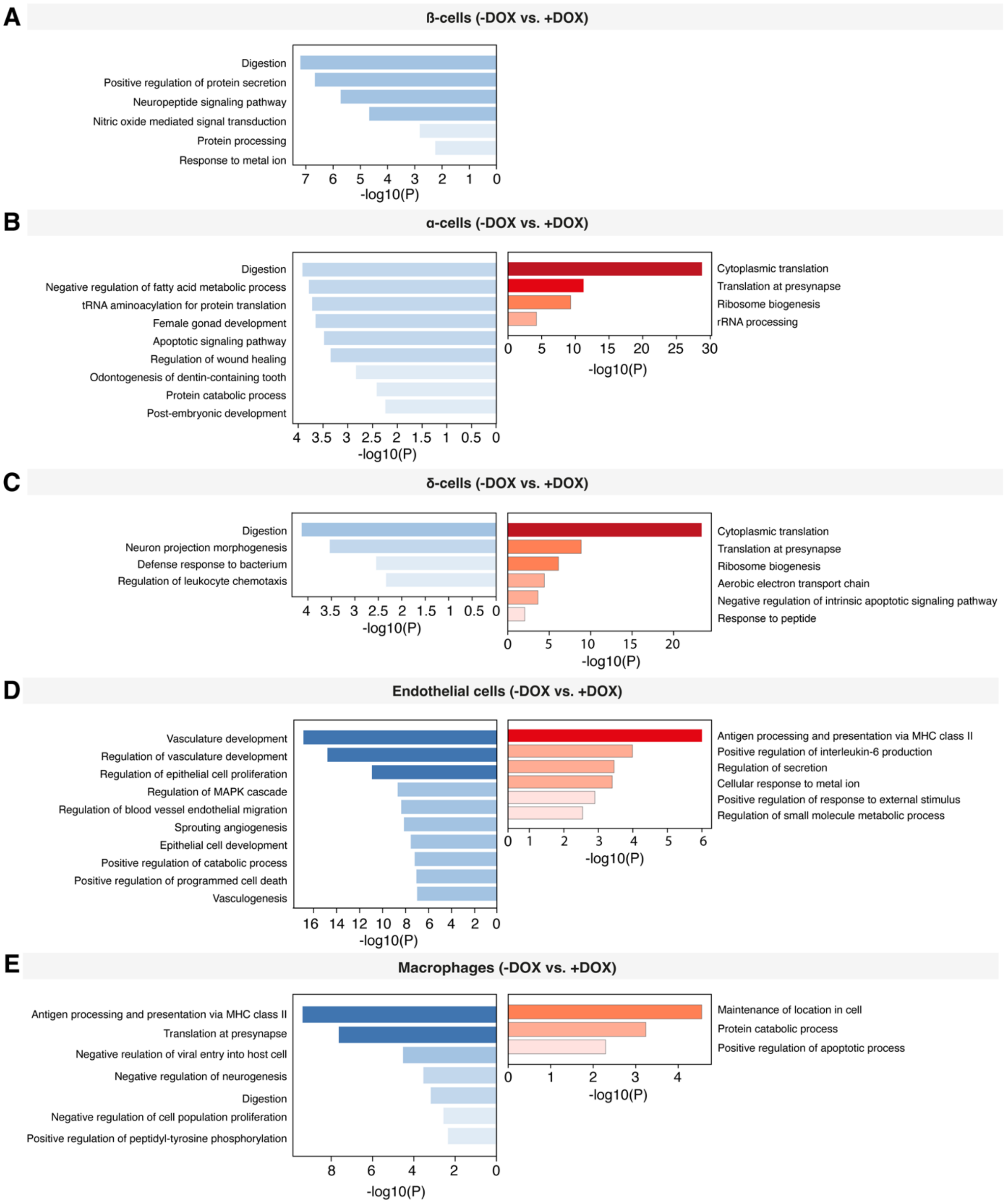

### Supplementary figure 4

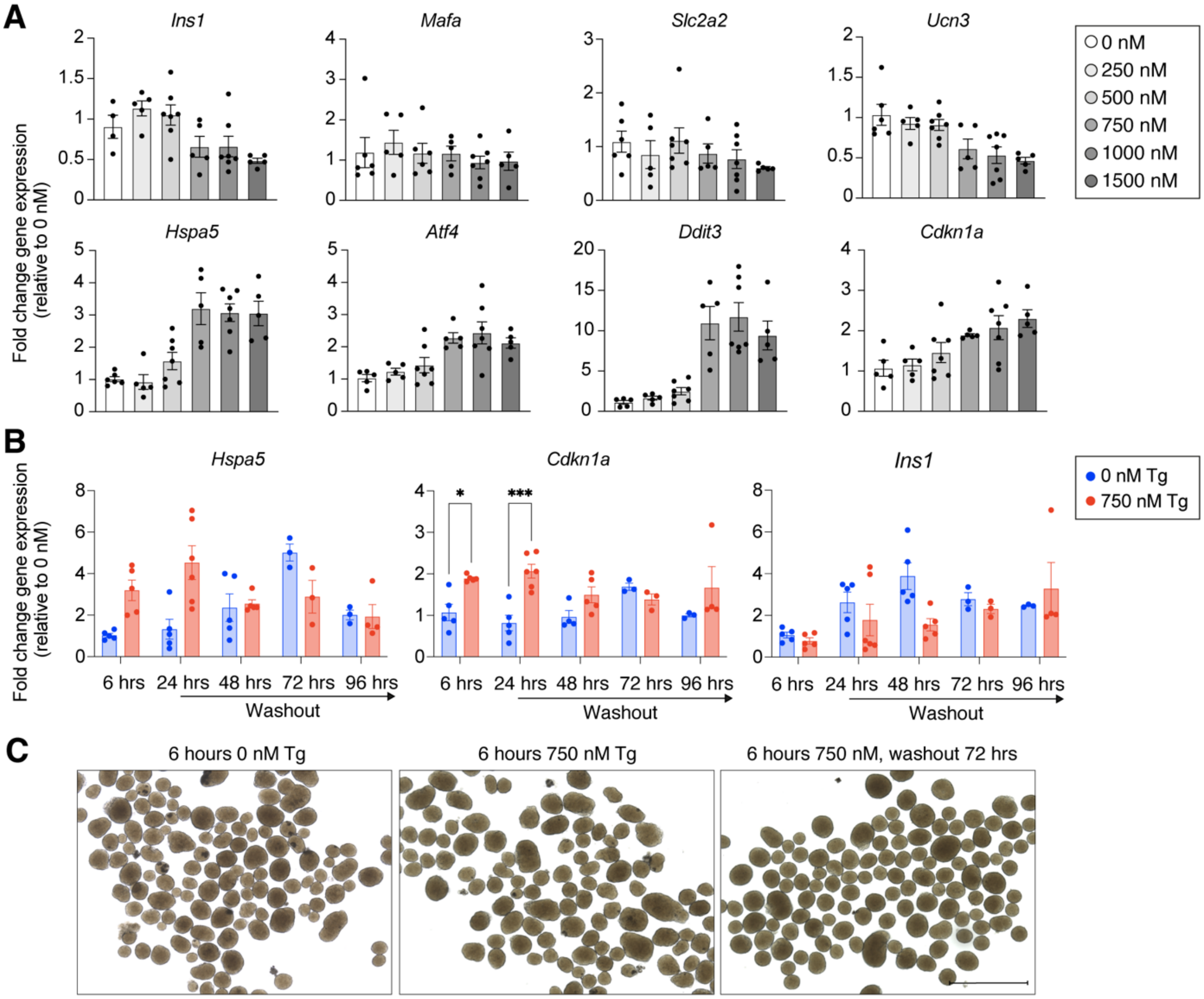
